# Tailoring cryo-electron microscopy grids by photo-micropatterning for in-cell structural studies

**DOI:** 10.1101/676189

**Authors:** Mauricio Toro-Nahuelpan, Ievgeniia Zagoriy, Fabrice Senger, Laurent Blanchoin, Manuel Théry, Julia Mahamid

## Abstract

Spatially-controlled cell adhesion on electron microscopy (EM) supports remains a bottleneck in specimen preparation for cellular cryo-electron tomography. Here, we describe contactless and mask-free photo-micropatterning of EM grids for site-specific deposition of extracellular matrix-related proteins. We attained refined cell positioning for micromachining by cryo-focused ion beam milling. Complex patterns generated predictable intracellular organization, allowing direct correlation between cell architecture and *in-cell* 3D-structural characterization of the underlying molecular machinery.

## Main

In parallel to the ongoing resolution revolution in cryo-electron microscopy for macromolecular structure determination^1^, cryo-electron tomography (ET) has matured to reveal the molecular sociology *in situ sensu stricto*^2-4^. Yet, cryo-ET of adherent mammalian cells can only be directly performed on their thin peripheries (< 300 nm). To reveal function-related structural variation, conformational states and function-related assemblies of macromolecules at the cell interior, thinning by advanced cryo-focused ion beam (FIB) has proved an optimal, artifact-free preparation method^2,5,6^. Specimen preparation for cellular cryo-ET, whether performed directly on thin cellular peripheries or following cryo-FIB micromachining, involves seeding of adherent cells directly on EM grids made of a biocompatible metal, e.g. gold. Standard EM grids are 3 mm diameter metal meshes overlaid with a delicate perforated thin film. Cells are typically allowed to spread, subjected to genetic or molecular perturbation to represent different physiological settings to be examined in molecular detail, that are then arrested by vitrification^7^. For cells to be thinned by cryo-FIB, they must be positioned roughly at the center of an individual grid square (Fig. 1a)^6^, within a few squares away from the grid center. Whilst the first requirement is necessary to allow access to cellular material for ablation by the FIB, the latter is posed by the subsequent requirement of stage tilt in the transmission electron microscope (TEM) for collection of tomographic tilt-series. Currently, optimization of grids for cellular cryo-ET is only carried out by adjusting the concentration of the cell suspension during seeding. However, adherent cells settle and adhere randomly on grids, often in the vicinity or directly on the metal grid bars, making them inaccessible to the FIB or to cryo-ET (Fig. 1a). Furthermore, presence of large clusters of cells increases the chance of incomplete vitrification due to slower heat transfer from the larger mass of material, and thus deteriorate structural preservation. Such technical hurdles have made the application of state-of-the-art cellular cryo-ET cumbersome and challenging. They further limit the advance of technical developments towards automation and high-throughput sample preparation. Here, we combined cellular cryo-ET with another technology in the fields of cell biology and biophysics, that of spatially controlled cellular environments. By adaptation of photo-micropatterning routinely applied to centimeter-scale glass slides for light microscopy-based assays^7,8^, we developed functionalized EM supports for directing cell positioning at high spatial accuracy, which ultimately renders molecular-resolution imaging of frozen-hydrated specimens more easily attainable.

**Figure 1.**
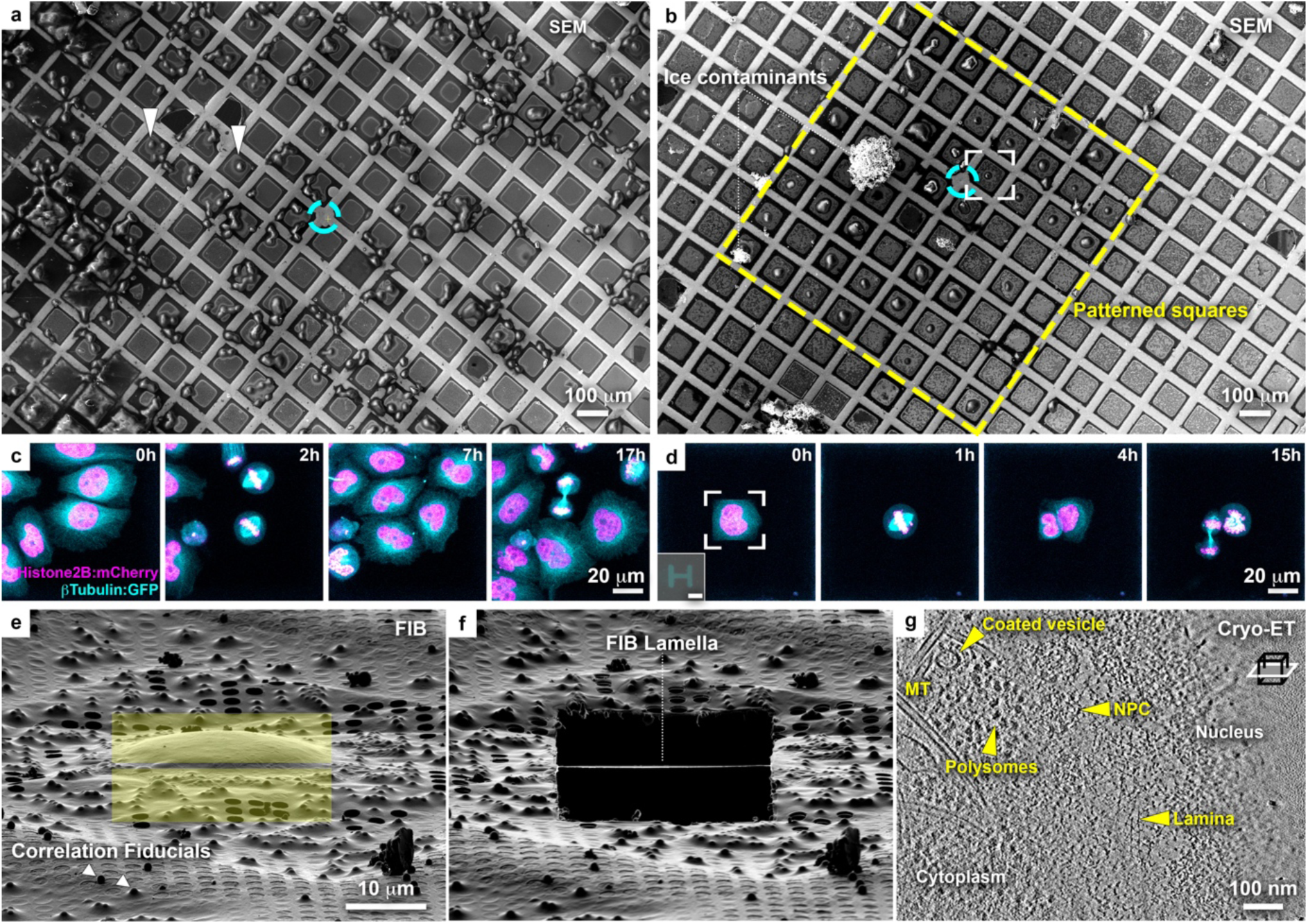
Micropatterning of cryo-EM grids refines preparation for cryo-FIB lamella micromachining from adherent mammalian cells. **(a)** Cryo-scanning electron micrograph (SEM) of HeLa cells grown overnight on a standard gold-mesh grid with SiO_2_ (R2/1) holey film. Cyan circle indicates grid center. Only a small fraction of the cells is optimally positioned for FIB-lamellae preparation (arrowheads). **(b)** HeLa cells grown overnight on a gold-mesh holey grid with 20 µm diameter disk patterns on 8 × 8 grid squares (yellow rectangle) around the grid center (cyan circle) treated with fibronectin. **(c-d)** HeLa cells, expressing GFP-tagged β-tubulin (Cyan) and mCherry-tagged histone (H2B-mCherry: magenta), seeded on a **(c)** control and **(d)** patterned gold-mesh grids with SiO_2_ (R1/20) holey film. Inset: H-shaped pattern induced the square cell shape. Scale: 20 µm. Cell-cycle was synchronized with a single Thymidine block (16 h), released into fresh medium (8h), followed by overnight live-cell imaging. A field of view of a single grid square is shown for both (c) and (d). Cells continue dividing ∼40 h post-seeding. **(e)** FIB shallow angle view on cell framed in (b). Yellow rectangles indicate patterns for milling. **(f)** Final lamella produced from cell in (e). **(g)** Tomographic slice, 6.8 nm thickness, of the nuclear periphery of the cell in (e). Lamella thickness determined from the tomographic reconstruction was 90 nm. NPC: nuclear pore complex; MT: microtubule.

We employed contactless and mask-free photo-micropatterning (Supplementary Fig. 1), ensuring preservation of the delicate support film and precise pattern generation within each grid square. The grids were plasma cleaned, rendering them hydrophilic, and coated with an anti-fouling biologically repelling agent (Polyethylene glycol (PEG)-brush, Supplementary Fig. 1a). Successively, specific areas were ablated with a UV-laser, either using a UV-pulsing laser scanned through the grid film on a standard confocal microscope (Supplementary Fig. 1b, Supplementary Fig. 2a-d) or continuous UV-illumination with a digital micro-mirror device (DMD) and a photo-activator (PLPP)^9^ (Supplementary Fig. 1b’). The DMD provides efficient area coverage of the grid film of approximately 150,000 µm^2^, equivalent to about 3 × 2 grid squares on a 200-mesh grid, and allows creating multiple, complex, and precisely positioned PEG-oxidized areas in ∼30 s at a resolution of 1.5 µm. The total patterned area can be extended by iterative montages of DMD expositions (Supplementary Fig. 3a). Following passivation and patterning, grids can be stored up to 30 days under hydrated conditions at 4°C. Grids were then functionalized with proteins that facilitated cell adhesion, and can be tailored to support growth and differentiation of various cell types.

**Figure 2.**
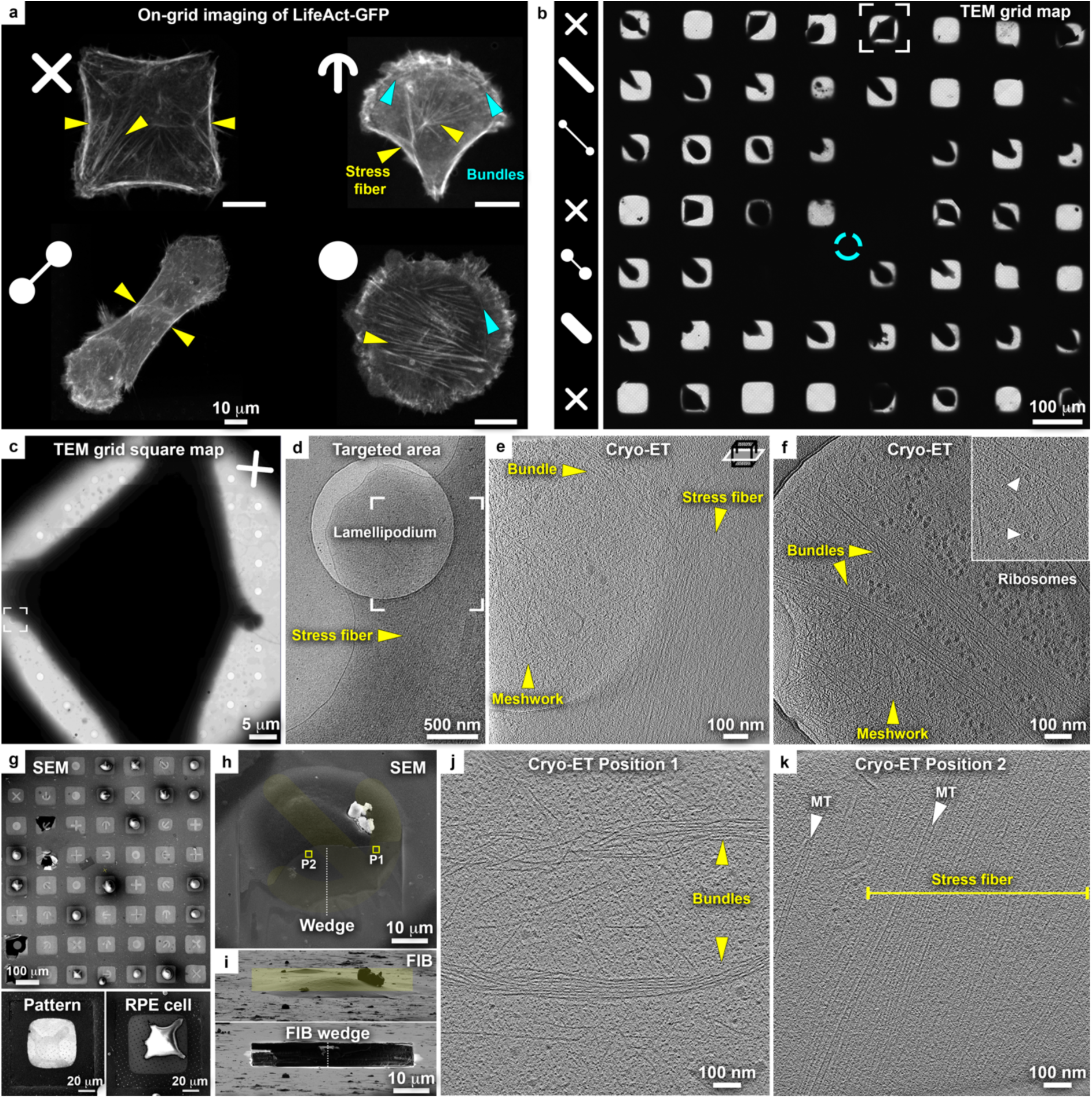
Cryo-EM grid micropatterning tailored for controlling cellular morphology and cytoskeletal architecture. **(a)** On-grid live-cell confocal microscopy of the actin organization in RPE1 LifeAct-GFP cells grown on complex micropatterns (gold-mesh, SiO_2_ film R1/4). Positioning of actin stress fibers (yellow arrowheads) correlate with the distinct patterns. Blue arrowhead: actin rings composed of putative bundles. **(b-f)** Cellular cryo-ET of RPE1 cells in peripheral thin regions. **(b)** Cryo-TEM map of grid with 8 × 7 patterned grid squares. Patterns per row are indicated (left column). Cyan circle: grid center. Framed cell is enlarged in (c). **(c)** Cryo-TEM micrograph of a grid square of the indicated cell in (b) grown on a cross-shaped pattern (rotated 90° counterclockwise from b). **(d)** Magnified cryo-TEM micrograph of the framed area in (c) (rotated 90° clockwise from c), targeted for tomography. **(e)** Tomographic slice of the specified area in (d), 6.8 nm thickness, showing the organization of actin filaments into a stress fiber and an isotropic meshwork in the adjacent lamellipodium. For full tomogram, see Video 2. **(f)** Tomographic slice of the periphery of another cell grown on an oval-shape pattern depicting actin meshwork, bundles, and unidentified hexameric macromolecular complexes in the vicinity of the basal and apical cell membranes (inset: arrowhead). For full tomogram, see Video 3. **(g-m)** Cellular thinning by cryo-FIB followed by cryo-ET. **(g)** SEM of RPE1 cells on a patterned titanium-mesh (SiO_2_ film R1/20) grid and overlaid with an image of the patterns. Bottom left: 2 keV SEM image of a cross-shaped micropatterned grid square. Right: SEM of RPE1 cell spreading on a cross-shaped pattern. **(h)** SEM of a cell grown on a crossbow-shaped pattern (yellow) overlaid with the SEM micrograph of a wedge (top view) produced by cryo-FIB milling. Squares indicate the positions of tomographic slices in (j) and (k). **(i)** Upper panel: FIB shallow angle view on cell in (h). Yellow rectangle indicates the pattern for milling. A thin wedge at the basal cell membrane is produced by ablating the top of the cell. Lower panel: cell after milling. **(j-k)** Tomographic slices of the positions 1 and 2 indicated in (h). Actin bundles likely equivalent to actin transverse arcs (parallel to but distant from the cell edge) and internal stress fibers -indicated in (a)-are found in locations expected according to the actin map in a crossbow-shaped RPE1 cell.

HeLa cells are a prominent model system in cell biology, and must be thinned to reveal structures positioned deep in their interior by cryo-ET. HeLa cells were seeded on fibronectin-functionalized micropatterned (30 µm disk-shape) grids. Reproducible seeding of single or double cells at the center of individual grid squares was achieved (Fig. 1b, Supplementary Fig. 2e-g, Supplementary Fig. 3b). Cells viability on micropatterned grids was confirmed by live-cell imaging: time-lapse light microscopy of cell-cycle synchronized cultures showed that cells spread and continue to divide 40h post-seeding (Fig. 1c-d, Supplementary Video 1). In this case, an H-shape pattern induced cells to adopt a rectangular appearance and can potentially be employed to produce defined geometries of cell division and of the mitotic spindle for structural analysis^10,11^. The majority of cells seeded on such grids were directly accessible for FIB thinning (Fig. 1b, e), highlighting the potential of grid micropatterning in streamlining challenging thinning techniques. Next, electron-transparent lamellae were generated (Fig. 1f, Supplementary Fig. 2g-h)^5^, and cryo-ET on the lamellae produced 3D-tomographic volumes of the nuclear periphery capturing previously-described molecular detail^4^ (Fig. 1g). Thus, the developed EM grid micropatterning method significantly contributes to optimization of advanced cellular cryo-ET pipelines, which encompass (i) vitrification, (ii) cryo-correlative light microscopy, (iii) micromachining by cryo-FIB milling, and (iv) cryo-ET.

Next, we generated complex patterns to control cell shape. Micropatterning on glass surfaces has been previously shown to induce well-defined cytoskeletal architectures and, as a result, a stereotypical internal organization of cellular organelles^7,8^. Here, we describe the actin network in Retinal pigment epithelium cells (RPE1) as a case of study and as a direct readout of the cellular response to adhesion on complex patterns. Tailored micropatterns induced reproducible cell morphology on grids (Fig. 2a)^12^. Each pattern elicited a distinguishable actin 3D architecture^8,13,14^, with spatially distinct peripheral and internal stress fibers, transverse arcs and isotropic branched meshworks. To elucidate the actin organization underlying these architectures, we first performed cellular tomography directly on the thin peripheries of micropatterned-shaped cells (Fig. 2b, Supplementary Fig. 4). A cell grown on a cross-shaped micropattern (Fig. 2c) was chosen aiming to target peripheral actin stress fibers as indicated by the live-cell actin map (Fig. 2a, upper-left). Cryo-TEM of the selected region exhibited a lamellipodium and a noticeable stress fiber (Fig. 2d, Supplementary Fig. 5a-d. See Supplementary Fig. 5e-k for additional examined cells). Cryo-ET of the targeted area revealed the presence of a peripheral bundle of aligned actin filaments comprising a stress fiber and the meshwork of the lamellipodium (Fig. 2e, Supplementary Video 2). Cryo-ET on a different, natively thinner, cell periphery displayed a meshwork of individual actin filaments, exhibiting the expected helical structure, and unidentified hexameric macromolecular complexes in the vicinity of the basal and apical cell membranes (Fig. 2f, Supplementary Fig. 5l-m, Supplementary Video 3).

To explore the organization of the cytoskeleton further away from the cell peripheries and to characterize spatially-predictable structures according to live-cell actin maps (Supplementary Fig. 6a-b), we further applied cryo-FIB milling to the micropattern-adherent RPE1 cells. Cryo-scanning electron microscopy (cryo-SEM) of a patterned grid shows the accurate single cell positioning at the centers of individual grid squares (Fig. 2g). Multiple different patterns can be generated on the same grid for direct comparison of different cellular architectures. The patterns’ design can be easily identified by direct imaging with low-voltage SEM or the micropattern-adopted cell shape (Fig. 2g, top and bottom panels), facilitating the overlay of the pattern design (Fig. 2g, top). An adherent RPE1 cell on a crossbow-shape micropattern was selected for thinning by means of cryo-FIB micromachining (Fig. 2h). The majority of the cell material was removed using a single rectangular area (Fig. 2i). This ablates the top of the cell, generating a thin wedge at the basal cell membrane^6^. Cryo-ET at positions 1 and 2, indicated on the wedge (Fig. 2h), displayed a curved transverse arc and a stress fiber, respectively (Fig. 2j-k, Supplementary Fig. 6c-d, Supplementary Video 4). Microtubules were seen lining the stress fiber, and fully embedded within this ultrastructure (Fig. 2k). These structures matched the expected actin organization in the crossbow pattern^13^, in which the observed bundles likely form part of the transverse arcs and the microtubules are aligned with the stress fiber at the basal part of the cell (Fig. 2a, upper-right). Such results provide an opportunity to uncover novel insights into cellular architecture following controlled cell shaping. Further data processing by semi-automated segmentation of tomographic volumes can deliver quantitative 3D structural information, including angular distributions of individual filaments and inter-filament distances in different cytoskeletal architectures, a level of structural information that is uniquely attainable by cryo-ET^15^. We further envision that the reproducible internal organization, encompassing positioning of cytoskeletal elements and stereotypical organelles organization in response to predetermined external physical cues posed by the micropatterns (Supplementary Fig. 6e)^13^, will assist in targeting specific internal elements for structural studies without resorting to challenging correlative approaches under cryogenic conditions.

In conclusion, photo-micropatterning of EM grids contributes to the advancement of refined, routine and user-friendly specimen preparations for *in-cell* structural biology. It further aids in solving technical challenges that have, thus far, hindered high-throughput FIB thinning preparations. This method will be instrumental for potential automation of the cryo-FIB milling process, deeply impacting the streamlining of cellular cryo-EM pipelines. This approach offers a unique opportunity to generate *in-cell* integrated insight into the structure and dynamics of macromolecules at nanometer-scale, broadening the scope of questions that can be addressed by state-of-the-art structural biology methods and bridging to other life-science disciplines.

## Acknowledgements

We are grateful to the ALMF and EMCF facilities at EMBL, especially to Stefan Terjung, Sebastian Schnorrenberg, Wim Hagen and Felix Weis, and to Giorgia Celetti for the kind gift of purified GFP protein. The Alvéole Lab company members Mehmet Akyuz, Grégoire Peyret, Laurent Siquier, Louise Bonnemay are acknowledged for advice and helpful discussions.

## Author contributions

M.T.-N., L.B., M.T., and J.M. conceived the study. M.T.-N., M.T. and J.M. design the experiments. M.T.-N., I.Z., F.S. and J.M. performed experiments. M.T.-N. and J.M. made the figures and wrote the manuscript, with contributions from all authors. M.T. and J.M. supervised the work.

## Data availability

The authors declare that the data supporting the findings of this study are available within the paper and its supplementary information files. The raw data of this study are available from the corresponding author upon reasonable request.

## Methods

### Cell lines and culture

Wild type HeLa Kyoto cells, and a double tagged line expressing both green fluorescent protein (GFP)-tagged β-tubulin from a bacterial artificial chromosome (BAC) and mCherry tagged histone from a plasmid construct (H2B-mCherry), were kindly provided by the Hyman Lab, Max Planck institute of Molecular Cell Biology and Genetics. HeLa cells were cultured in Dulbecco’s modified Eagle’s medium (DMEM GlutaMAX; ThermoFischer Scientific, Schwerte, Germany), while RPE1 (Retinal Pigment Epithelial human cells) expressing LifeAct-GFP^16^ were cultured in DMEM F-12 GlutaMAX. Cells were incubated at 37°C with 5 % CO_2_, and supplemented with 10 % (v/v) fetal bovine serum (FBS), 100 mg/mL penicillin, 100 mg/mL streptomycin. A 0.5 mg/mL geneticin (G418) for the BAC-tagged lines and Puromycin (1 µg/ml) for cells carrying the plasmids were used. FluoroBrite DMEM supplemented with 2mM L-glutamine (ThermoFischer Scientific, Schwerte, Germany) was used for live cell fluorescence imaging.

### Electron microscopy grids

Gold (Au) or Titanium (Ti) 200-mesh grids with a holey 12 nm thick SiO_2_ film, either R2/1 (2 μm holes separated by 1 μm spacing), R1/4 or R1/20 (Quantifoil Micro Tools, Jena, Germany) were employed in this study. Titanium-mesh grids, and SiO_2_ films replacing the commonly used amorphous carbon Quantifoil, provided stiffer and more robust supports for the multiple grid processing, and cell culture steps described in the methods. Both titanium (Fig. 2g-h) and SiO_2_ film were demonstrated to be biocompatible as observed by live-cell imaging (Fig. 1c-d). The commonly used amorphous carbon Quantifoil films were also compatible with the photo-micropatterning method, yet require sputtering of an additional carbon layer on the support^4,5^ for easier handling during cell culture.

Increased amount of film over holes promoted better cell adhesion. R2/1 and R1/4 films were advantageous for direct tomography of peripheral cellular areas, while R1/20 and R1/4 films were more suitable for cellular thinning by cryo-FIB milling as the majority of the film was removed during thinning.

### Grid passivation

#### One-step

Grids were oxidized and rendered hydrophilic using a low-pressure Diener Femto plasma cleaner. Grids were placed onto a glass slide and both sides were plasma cleaned at 100W power with a 10 cm^3^/min flow rate of oxygen gas for 30-40 s. Next, grids were incubated on droplets of poly-L-lysine grafted with polyethylene glycol (PLL(20)-g[3.5]-PEG(5), SuSoS AG, Dübendorf, Switzerland) at a concentration of 0.5 mg/ml in 10 mM Hepes pH 7.4, for 1h at room temperature or overnight at 4°C, on parafilm in a humid chamber (parafilm sealed dish with soaked filter paper). Following passivation, the grids were blotted with filter paper from the back, allowed to dry for few seconds, and immediately placed on a drop of the next micropatterning solution (depending on the micropatterning method) in a sealed glass bottom ibidi µ-Dish 35 mm low (ibidi, Martinsried, Germany).

#### Two-step

As an alternative, a two-step treatment of the grids was also tested. First, grids were plasma cleaned and then incubated on droplets of 0.01 % PLL (Sigma Aldrich, St. Louis, MO) on parafilm in a humid chamber overnight at room temperature. Next, the grids were rinsed five times in 100 mM NaHCO_3_ pH 8.4, and incubated for 1-2 h with 50 mg/ml PEG-sva dissolved in the same solution (Laysan Bio, Arab, US). Subsequently, the grids were washed five times in 100 mM NaHCO_3_ pH 8.4. This treatment can be potentially applied in the absence of a plasma cleaner.

Both protocols were successful for grid passivation followed by photo-micropatterning, as judged from fluorescence light microscopy imaging of GFP-adsorption that was restricted to the PEG-free patterns. However, a one-step PLL-g-PEG passivation was preferred for time consideration.

### Micropattern design

Micropatterns were designed in Inkscape (http://www.inkscape.org/) as 8-bit binary files and exported as png files, which can be loaded into the Leonardo software v4.12 (Alvéole Lab, Paris, France).

### Micropatterning and functionalization of grids

#### Nanoablation by a 355 nm pulse laser

An inverted confocal Olympus FluoView 1200 (Olympus, Hamburg, Germany) microscope was used, equipped with a UV pulsed laser source of 355 nm (PNV-001525-140, Teem Photonics, Meylan, France), a UPLSAPO 63x (NA 1.35) oil objective, and a standard PMT or GaAsP PMT detectors. The 355 nm laser had an average power of 50 mW, 300 ps pulse width, 1kHz repetition rate, and a maximum energy per pulse of 20 µJ. After passivation with PLL-g-PEG, grids were blot-dried from the back with a filter paper and quickly placed with the SiO_2_ film facing down (towards the objective) on 1-3 µl of Hepes 10 mM pH 7.4 or PBS in a sealed glass bottom ibidi µ-Dish 35 mm low (ibidi, Martinsried, Germany). High humidity was kept using water-soaked filter paper inside the dish to avoid solution evaporation during micropatterning. Transmission and reflection were observed with a 488 nm laser. The patterns (ROIs) were of square or circular shape (20, 30 or 40 µm diameter) generated using the Olympus FV 10-ASW software v04.02.03.02. Photo-micropatterning was performed using 10-11 % laser power, 40 μs per pixel and 10 iterations. Individual grid squares were targeted at a time, the film focused and the laser applied. Micropatterning of a 4 × 4 grid squares area (200-mesh grid: ∼260,000 µm^2^) took approximately 8 min. Potentially, a lower magnification objective can be used in order to pattern more grid squares at the same time in order to optimize patterning, provided that the film is flat and at even height to maintain all areas in the focus plane. Titanium grids had a consistent film flatness aiding quick focusing on each grid square, facilitating the micropatterning using this technique.

#### PRIMO™ (DMD-based illumination + Photo-activator)

An inverted Nikon microscope Ti-E equipped with a CFI Super Plan FLuor 20x ELWD (NA 0.45) lens with high UV-transmission, a Perfect Focus System 3, an ORCA-Flash 4.0 LT CMOS camera (Hamamatsu, Japan), a motorized stage (Märzhäuser, Wetzlar, Germany), and the Primo™ micropatterning module (Alvéole Lab, Paris, France) was used. Grid micropatterning was performed using digital mirror device (DMD) to generate a spatially controlled laser illumination of the sample (Primo™, Alvéole Lab, Paris, France), with a resolution limit of ∼1.2 µm. After passivation with PLL-g-PEG, grids were blot-dried from the back with a filter paper and quickly placed on a 1-3 µl of PLPP (4-benzoylbenzyl-trimethylammonium chloride, 14.5 mg/ml) drop in a sealed glass bottom ibidi µ-Dish 35 mm low (ibidi, Martinsried, Germany). High humidity was kept using water-soaked filter paper inside the dish to avoid PLPP evaporation. The dish, with 1-4 grids at a time, was placed on the microscope stage and photo-patterning was controlled with the µmanager software v1.4.22 by the Leonardo plugin software v4.12 (Alvéole Lab, Paris, France) using the stitching mode and a 375 nm (4.5 mW) laser, applying a dose of 800-1000 mJ/mm^2^ equivalent to ∼30 s per DMD exposition. Micropatterning of 8 × 7 grid squares area (200-mesh grid: ∼900,000 µm^2^) took 3-7 min depending on the total dose and grid positioning with respect to the DMD mirror illumination. Grids were promptly retrieved from the PLPP solution, washed in a 300 µl drop of water, and two consecutive washes in 300 µl drops of PBS. Patterned one-step passivated grids were stored wet in PBS at 4°C in a humid chamber, remaining functional for at least 30 days.

#### Grid functionalization

For functionalization following PEG ablation in micropatterns using both methods, grids were incubated at room temperature for 1h in a 20 µl drop of either 100 µg/ml of GFP, 50 µg/ml fibronectin (ThermoFischer Scientific, Schwerte, Germany), or 30 µg/ml of fibrinogen-Alexa546 (ThermoFischer Scientific, Schwerte, Germany) on parafilm and, subsequently, washed 3 times in 300 µl drops of PBS. Fibrinogen was prepared in 100 mM NaHCO_3_ pH 8.4, hence washes were performed in the same buffer. Grids incubated with fibronectin remained functional for at least 10 days in PBS at 4°C in a humid chamber. The maximum active life time of the micropatterned grids as well as protein functionalization remain unknown, as this will also depend on the protein stability itself.

All patterning steps and grid treatments were performed under sterile conditions using a Bunsen burner. Grids were handled with a tweezer n° 55 (Dumont, Montignez, Switzerland).

#### Comparison between the two patterning approaches

Due to the pulsing nature of the laser in the first approach, it has to scan the region of interest in order to ablate the anti-fouling agent. The action of the laser leaves an impression on the film that is visible by light microscopy (Supplementary Fig. 2a-b). The engraving can be further observed in detail by FIB (side view) and SEM (top view) imaging of a grid square (Supplementary Fig. 2c-d). Importantly, the film engraving at the conditions used did not seem to cause a deterioration of the SiO_2_ layer. In fact, proper cell adhesion to the fibronectin-coated patterns (Supplementary Fig. 2e-f) and the subsequent vitrification of grids appeared unaltered (Supplementary Fig. 2g). Furthermore, followed by cell settling on the patterns, they strictly adhere and divide on the micropatterned regions (Supplementary Fig. 2f), denoting cell viability and the effectiveness of the passivation. Some agglomeration of cells can be appreciated in the Supplementary Fig. 2e. This can be overcome by optimizing cells seeding conditions using fewer cells/cm^2^ and less time of seeding (earlier transfer to a new cell-free dish) along with use of a cell strainer. Using this method, a directed spatial positioning of cells was attained, which was valuable for cryo-FIB milling (Supplementary Fig. 2g-j), demonstrating and further supporting the advantages of micropatterning for the cellular cryo-ET pipeline optimization.

##### Timing

while the Primo device takes 30 s per DMD run (for a dose of 1000 mJ/mm^2^ and covering a 3 × 2 grid squares on a 200-mesh grid), the 355 nm-pulse laser patterning takes ∼10-15 s per grid square considering a disk-shaped pattern of 20-30 µm diameter. A user familiar with the 355 nm laser technique can pattern a 4 × 4 grid square area in ∼8 min, while the Primo technology covers a similar area in ∼1 min.

##### Resolution

the 355 nm-pulse scanning laser can yield a much higher spatial lateral resolution limited by the light diffraction limit equivalent to ∼250 nm, in comparison to the Primo performance that is limited to ∼1.5 µm.

### Cell seeding

Non-patterned grids were plasma cleaned. Cells were detached from cell culture flasks using 0.05 % trypsin-EDTA and seeded on pre-treated (either patterned or non-patterned) Quantifoil grids in glass bottom ibidi µ-Dish 35 mm high (ibidi, Martinsried, Germany).

Cells were seeded on fibronectin micropatterned surfaces right after being passed through a cell 40 µm pore-size cell strainer (Corning, Amsterdam, Netherlands) at a density of 2× 10^4^cells/cm^2^ for HeLa and 8×10^3^cells/cm^2^ for RPE1 cell lines. After seeding, grids were incubated for 1.5-2 h for HeLa cells or 20-35 min for RPE1 cells. Next, grids were transferred to a new cell-free dish and incubated at 37°C with 5 % CO_2_ to allow cell adhesion to the grids. Transfer to a new dish was beneficial to remove cells that were non-specifically attached to areas outside the patterns. Cells were vitrified 4-6 h post-transfer for RPE1 cells (to attain a higher number of grid squares with a single cells) or after overnight incubation for HeLa cells. At least 50 grids have been seeded with either HeLa or RPE1 cells obtaining reproducible results with cells settling and adhering to the micropatterned areas.

### Live-cell confocal imaging

Cell-cycle synchronization aimed at validating cell viability after adhesion on micropatterned grids by live cell imaging. HeLa cells were allowed to adhere overnight (16 h) in the presence of 2 mM Thymidine (S-phase block; Merck, Darmstadt, Germany) and released into fresh medium. Imaging started 8h post-release and time-laps movies were recorded overnight.

Time lapse imaging of HeLa cells on grids (Fig. 1c, d) was performed in a Zeiss LSM 880 Airyscan microscope (Carl Zeiss, Jena, Germany) using confocal detectors and a Plan-Apochromat 20 × (NA 0.8) objective. Field of view: 425 × 425 µm^2^. Pixel size of 149 nm, and 1 µm as *z*-step (total z-depth: 32 µm). Pixel dwell time: 0.28 µs. Master gain 850 for both channels. GFP was detected using a 488 nm line of Argon laser with a 1.5 % power and a 490-550 nm bandpass emission filter. mCherry was detected using a 561 nm DPSS laser at a 0.15% power, and a bandpass emission filter of 570-660 nm. Time-lapse imaging was performed with 1h time resolution using the Zen Black 2.3 SP1 software v14.0.15.201 and MyPic VBA macro^17^. XZ images of the dish surface reflection was acquired and processed by the MyPic VBA macro to define axial position for *z*-stack acquisition at each time point. Each stack was post-processed in Fiji^18^. Briefly, channels were split, maximum intensity projections produced, channels combined into a single image per time point, and all time points combined into a movie stack.

### Zeiss Airyscan microscopy

AiryScan microscopy of RPE1 cells on patterned grids (Fig. 2a) was performed on a Zeiss LSM 880 AiryScan microscope (Carl Zeiss, Jena, Germany), using an AiryScan detector and a C-Apochromat 40x (NA 1.2) water immersion objective. Optimal sampling conditions for AiryScan acquisitions were achieved by selecting SR (super-resolution) scanning modality. Pixel size of 50 nm, and a 225 nm *z*-step (total z-depth: 10-15 µm). Pixel dwell time: 0.64-1.18 µs. Master gain: 850-900. LifeAct-GFP was detected using a 488 nm line of Argon laser with a power of 1.5-2 % and a 495-550 nm bandpass emission filter. Stack datasets were post-processed in Zen Black 2.3 SP1 software v14.0.15.201 (Zeiss) to combine the multiple Airyscan 32-detector array images into deconvolved final images with high SNR and resolution.

### Widefield microscopy imaging

Epifluorescence images (Fig. 2d inset, Supplementary Fig. 2) were recorded using a Zeiss Axio Observer.Z1 widefield microscope (Carl Zeiss, Jena, Germany) with an AxioCam MRm CCD camera, a A-Plan 10x (NA 0.25) Ph1 and a LD-Plan Neofluar 20x (NA 0.4) Ph2 Corr objectives. GFP was imaged with a 38 HE filter set and mCherry with a 43 HE filter set. Cells were imaged at 37° and 5 % CO_2_ with the Zen (blue edition) 2.3 software v2.3.69.1000. Images were processed histogram adjusted, and cropped in Fiji.

### Vitrification

Grids were blotted from the reverse side of the film and immediately plunged into a liquid ethane or ethane/propane mixture at liquid nitrogen temperature using a Leica EM GP plunger (Leica Microsystems, Vienna, Austria). The plunger was set to 37°C, 99 % humidity, and blot time of 2 s for R2/1, and 2.5 s for R1/4 and R1/20 grids. The frozen grids were stored in sealed boxes in liquid nitrogen until further processing.

### Cryo-scanning electron microscopy and focused ion beam milling

Cryo-FIB lamella preparations was performed as described in^5^, on a dedicated Aquilos dual-beam microscope equipped with a cryo-transfer system and a cryo-stage (ThermoFisher Scientific, Brno, Czech Republic). Plunge frozen grids were fixed into autogrids modified for FIB preparation (a generous gift from the Max Planck institute of Biochemistry, Martinsried, Germany), mounted into a shuttle (ThermoFisher Scientific, Brno, Czech Republic) and transferred into the dual-beam microscope via a load-lock system. During FIB operation, samples were kept at constant liquid nitrogen temperature using an open nitrogen-circuit, 360° rotatable cryo-stage. To improve sample conductivity and reduce curtaining artifacts during FIB milling, the samples were first sputter-coated with platinum *in situ* (10 mA, 20 s), and then coated with organometallic platinum using the *in situ* gas injection system (GIS, ThermoFisher Scientific, Eindhoven, Netherlands) operated at room temperature, 10.6 mm stage working distance and 7 s gas injection time. Appropriate positions for FIB preparations were identified and recorded in the MAPS 3.3 software (ThermoFisher Scientific, Brno, Czech republic), and the eucentric height refined per position. Lamellae or wedges were prepared using Gallium ion beam at 30 kV, stage tilt angles of 20° for lamellae and 12°-13° for wedges. Lamella or wedge preparations were conducted in a stepwise manner, starting 5μm away from the area of interest with currents of 1 nA, gradually reduced to a current of 50 pA for the final cleaning steps. Progress of the milling process was monitored using the scanning electron beam operated at 10 keV and 50 pA (or 2keV for visualization of micropatterns). For improved conductivity of the final lamella for specimens intended for phase plate tomography, we again sputter coated the grid after cryo-FIB preparation with platinum (10 mA, 3 s). Grid were stored in sealed boxes in liquid nitrogen until further processing.

### Cryo-electron tomography

Cryo-electron microscopy data were collected on a Titan Krios microscope operated at 300 kV (ThermoFisher Scientific, Eindhoven, Netherlands) equipped with a field-emission gun, a Quantum post-column energy filter (Gatan, Pleasanton, CA, USA), a K2 Summit direct detector camera (Gatan) and a Volta phase plate (ThermoFisher Scientific, Eindhoven, Netherlands). Data were recorded in dose-fractionation mode using acquisition procedures in SerialEM software v3.7.2^19^. Prior to the acquisition of tilt-series, montages of the grid squares or lamella were acquired at ∼2 nm/pix. Tilt-series using a dose symmetric scheme^20^ were collected in nano-probe mode, EFTEM magnification 42,000x corresponding to pixel size at the specimen level of 3.37 Å, 3-4 µm defocus, tilt increment 2° with constant dose for all tilts, total dose ∼ 120 e^-^/Å^2^. The pre-tilt of lamellae with respect to the grid plane due to cryo-FIB milling at shallow angles (10-15°) was corrected for by tilting the stage on the microscope. Conventional tilt-series, without Volta phase plate (VPP), were acquired with an objective aperture and a beam tilt of 4 mrad for autofocusing (tomograms in Fig. 2e and 2f; Supplementary Fig. 5). A fraction of the tomographic tilt-series was acquired with the VPP (tomograms in Fig. 1g; Fig. 2j, k). Alignment and operation of the Volta phase plate were essentially carried out as described previously, applying a beam tilt of 10 mrad for autofocusing^4^. For conventional defocus data, a total of 30 tomograms were acquired on the peripheral areas of 13 micropatterned cells (from all micropattern designs shown in Fig. 2b). A total of 31 VPP tomograms were acquired from 8 wedges, equivalent to 8 cells grown on crossbow-, cross-, dumbbell-, oval-, and disk-shape micropatterns.

### Data processing

Prior to tilt-series alignment, the projection movies were corrected for beam induced drift in the SerialEM plugin. Tilt series alignment and tomographic reconstructions were performed using the IMOD software package v4.9.0^19^. In absence of fiducial gold nanoparticles in the FIB-lamellae or wedges, alignment of tilt-series images was performed with patch-tracking. Final alignment of the tilt-series images was performed using the linear interpolation option in IMOD without CTF correction. Aligned images were binned to the final pixel size of 13.48 Å. For tomographic reconstruction, the radial filter options were left at their default values (cut off, 0.35; fall off, 0.05). Tomograms from Fig. 2e and Supplementary Fig. 5c-d were treated with an anisotropic nonlinear diffusion denoising algorithm implemented in IMOD to improve signal-to-noise ratio.

## References

1. Kühlbrandt, W. Science 343, 1443–1444 (2014). doi:10.1126/science.1251652

2. Pfeffer, S. & Mahamid, J. Curr Opin Struct Biol 52, 111–118 (2018). doi:10.1016/j.sbi.2018.08.009

3. Beck, M. & Baumeister, W. Trends Cell Biol 26, 825–837 (2016). doi:10.1016/j.tcb.2016.08.006

4. Mahamid, J. et al. Science 351, 969–972 (2016). doi:10.1126/science.aad8857

5. Schaffer, M. et al. J Struct Biol 197, 73–82 (2017). doi:10.1016/j.jsb.2016.07.010

6. Rigort, A. et al. Proc Natl Acad Sci U S A 109, 4449–4454 (2012). doi:10.1073/pnas.1201333109

7. Azioune, A., Carpi, N., Tseng, Q., Thery, M. & Piel, M. Methods Cell Biol 97, 133–146 (2010). doi:10.1016/S0091-679X(10)97008-8

8. Thery, M. J Cell Sci 123, 4201–4213 (2010). doi:10.1242/jcs.075150

9. Strale, P. O. et al. Adv Mater 28, 2024–2029 (2016). doi:10.1002/adma.201504154

10. Thery, M., Jimenez-Dalmaroni, A., Racine, V., Bornens, M. & Julicher, F. Nature 447, 493–496 (2007). doi:10.1038/nature05786

11. Thery, M. et al. Nat Cell Biol 7, 947–953 (2005). doi:10.1038/ncb1307

12. Engel, L. et al. Preprint at https://doi.org/10.1101/657072 (2019).

13. Thery, M. et al. Proc Natl Acad Sci U S A 103, 19771–19776 (2006). doi:10.1073/pnas.0609267103

14. Senger, F. et al. Preprint at http://dx.doi.org/10.1101/578799 (2019).

15. Jasnin, M. & Crevenna, A. H. Biophys J 110, 817–826 (2016). doi:10.1016/j.bpj.2015.07.053

16. Vignaud, T. et al. J Cell Sci 125, 2134–2140 (2012). doi:10.1242/jcs.104901

17. Politi, A. Z. et al. Nat Protoc 13, 1445–1464 (2018). doi:10.1038/nprot.2018.040

18. Schindelin, J. et al. Nat Methods 9, 676–682 (2012). doi:10.1038/nmeth.2019.

19. Kremer, J. R., Mastronarde, D. N. & McIntosh, J. R. J Struct Biol 116, 71–76 (1996). doi:http://dx.doi.org/10.1006/jsbi.1996.0013

20. Hagen, W. J. H., Wan, W. & Briggs, J. A. G. J Struct Biol 197, 191–198 (2017). doi:10.1016/j.jsb.2016.06.007

