## Supplementary_Information for "Tailoring cryo-electron microscopy grids by photo-micropatterning for in-cell structural studies"

### *Supplementary Figures*

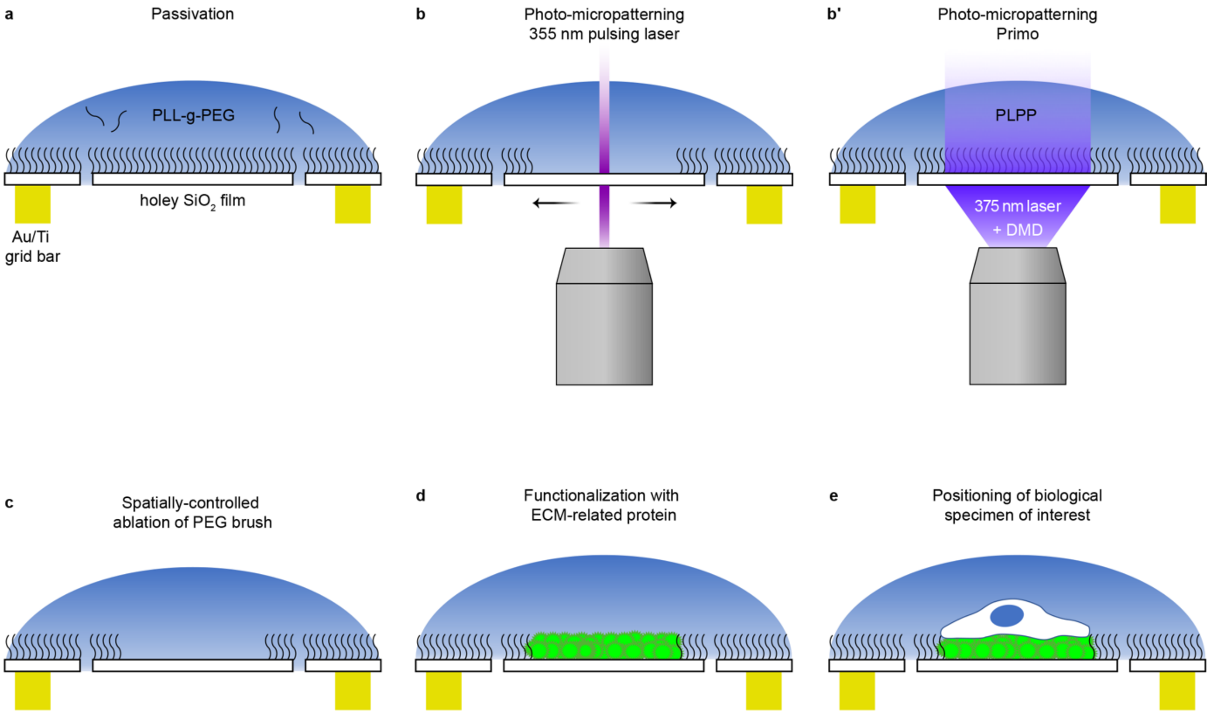


**Supplementary Figure 1. Micropatterning of electron microscopy grids for high spatial accuracy cell positioning.**

**(a)** Grid passivation with anti-fouling agent PLL-g-PEG generates an organized repulsive PEG brush at the surface. **(b)** UV laser application using a 355 nm pulsing laser scanned through the region of interest causes ablation of the passivation layer. **(b')** UV laser application using a digital-micromirror device (DMD) and the photo-initiator (PLPP) to locally oxidize the passivation layer. **(c)** Spatially constrained ablation of the PLL-g-PEG passivation layer. **(d)** Grid functionalization with extracellular matrix (ECM)-related proteins. **(e)** Cell seeding at the functionalized micropatterned areas.

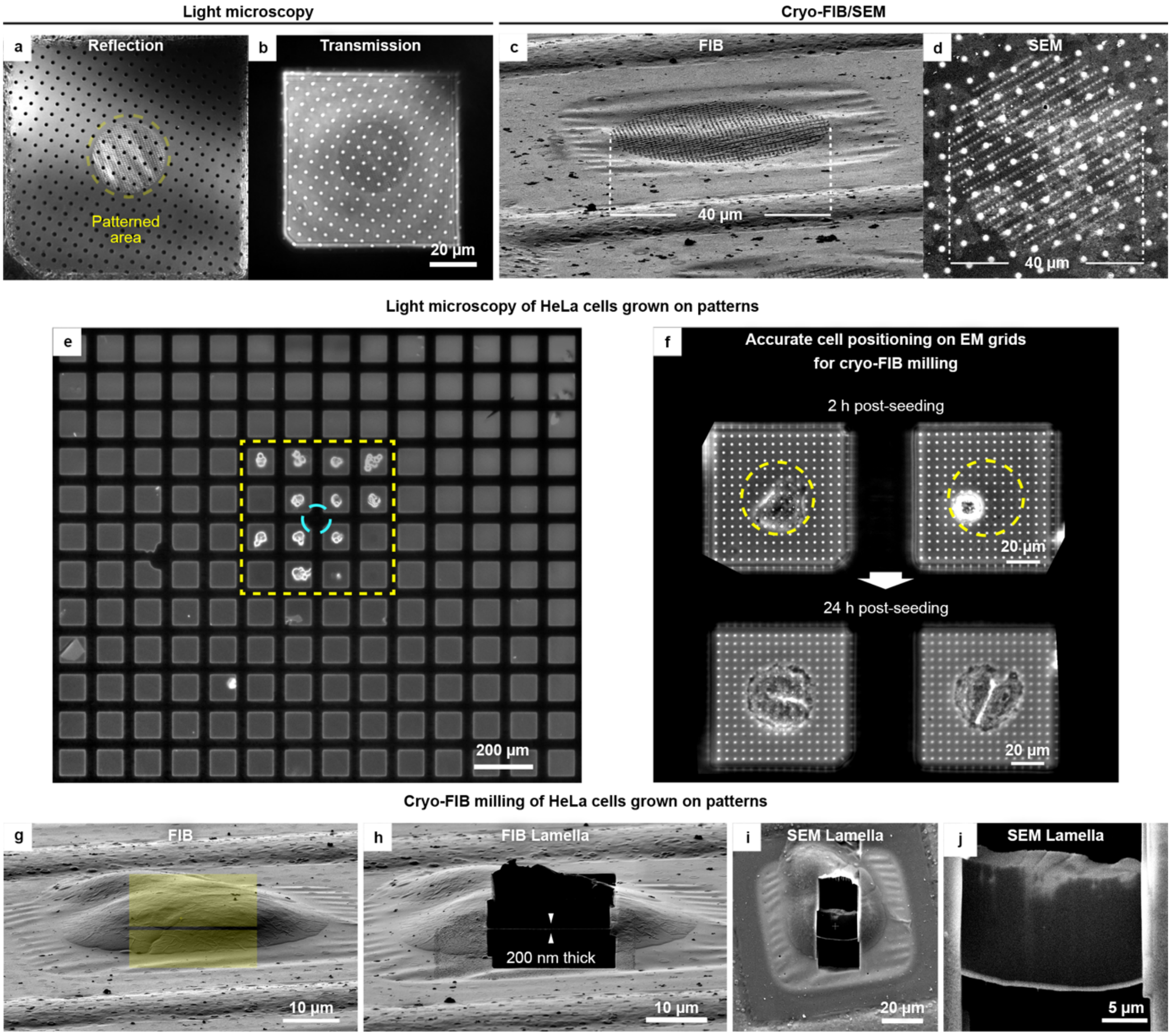


**Supplementary Figure 2. Photo-micropatterning of EM grids using a UV-355 nm pulsing laser.**
**(a-b)** Reflection and transmission light microscopy of micropatterns generated by a 355 nm wavelength pulsing laser scanned on the grid film to generate a 30 µm disk-shaped area (gold-mesh grid, $SiO_2$ film R1/4). The micropatterned area, imaged in the confocal microscope immediately post-laser application, can be identified by the impression left as a result of laser pulses on the $SiO_2$ film. **(c-d)** Cryo-FIB/SEM imaging of a micropatterned grid post-vitrification displaying the engraving made by the laser. **(e)** Gold-mesh holey grid micropatterned 4 x 4 grid squares (30 µm disk-shape) around the grid center (cyan circle), treated with fibronectin and seeded with HeLa cells. Cells are constrained to the patterned area. **(f)** Light microscopy imaging of a grid 2.5 h after seeding (upper-panel), displaying single cells at the micropatterned circular region. Cell division (24 h post-seeding, lower-panel) is restricted to the micropatterned area. **(g)** FIB shallow angle view of a cell grown on a disk-shaped pattern under cryogenic conditions. Yellow rectangles indicate the pattern for milling to produce a thin lamella through the cell center. **(h)** FIB view of the cell after milling to generate a ~200 nm thin lamella. **(i)** SEM top view, and zoom-in **(j)**, of the lamella from (h).

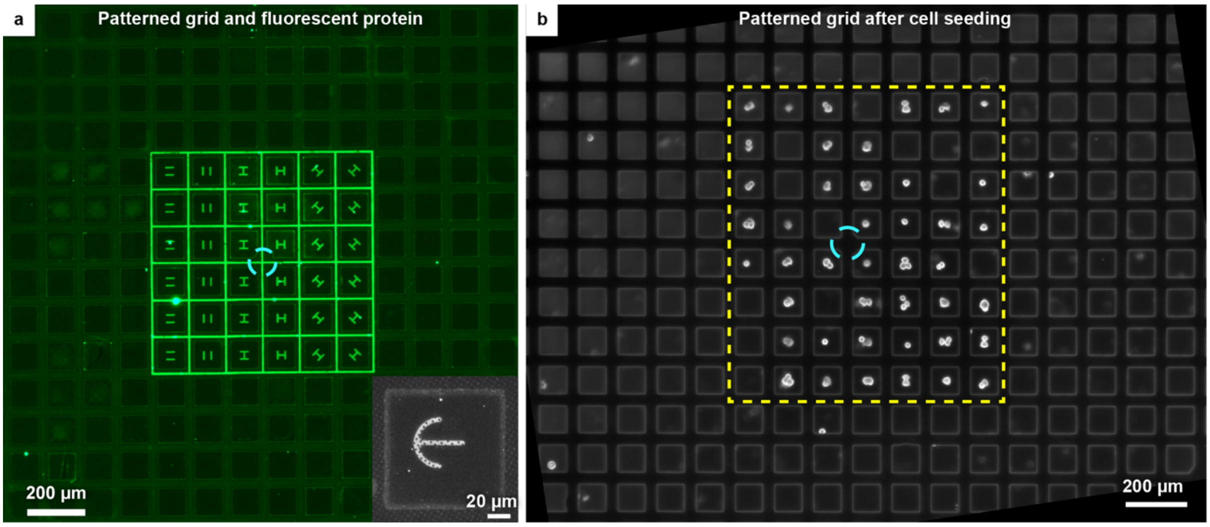

**Supplementary Figure 3. Photo-micropatterning of EM grids using a digital-micromirror device and the photo-initiator PLPP.**

**(a)** Gold-mesh holey ($SiO_2$ film) grid with micropatterned 6 x 6 grid squares (H-shape of 30 µm size) around the grid center (cyan circle) using the DMD plus PLPP method and on-step passivation. The grid was incubated with GFP protein to validate micropatterning. Grid bars areas are also patterned (3µm lines). Inset: cross-bow pattern generated following two-step passivation coated with fibrinogen-Alexa546 to validate micropatterning. **(b)** Gold-mesh holey ($SiO_2$ film) grid with micropatterned 8 x 7 grid squares (H-shape of 30 µm size) around the grid center (cyan circle) using the DMD plus PLPP method, treated with fibronectin, seeded with HeLa cells and imaged at 24h post-seeding.

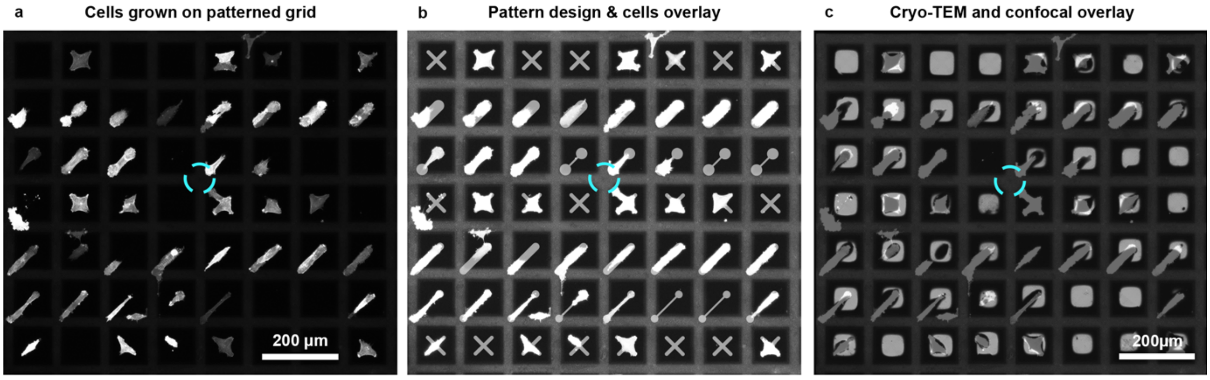

**Supplementary Figure 4. Correlation of grid maps from fluorescence and transmission electron microscopy.**

**(a)** On-grid live-cell imaging (confocal microscopy) to generate a grid map of RPE1 LifeAct-GFP culture (4h post-seeding) on gold-mesh ($SiO_2$ film R1/4) grid with 8 x 7 patterned grid squares. Micrograph is a maximum intensity projection of a z-stack. **(b)** Fluorescence microscopy map from (a) overlaid with the pattern design. The micrograph is contrast adjusted for better visualization. **(c)** Cryo-TEM map of same grid (vitrified ~1h after live-cell imaging), and overlaid with the fluorescence microscopy map. Cells are dark in the TEM map and light in the light microscopy map. Cyan circle: grid center. Micrographs are flipped relative to Fig. 2b.

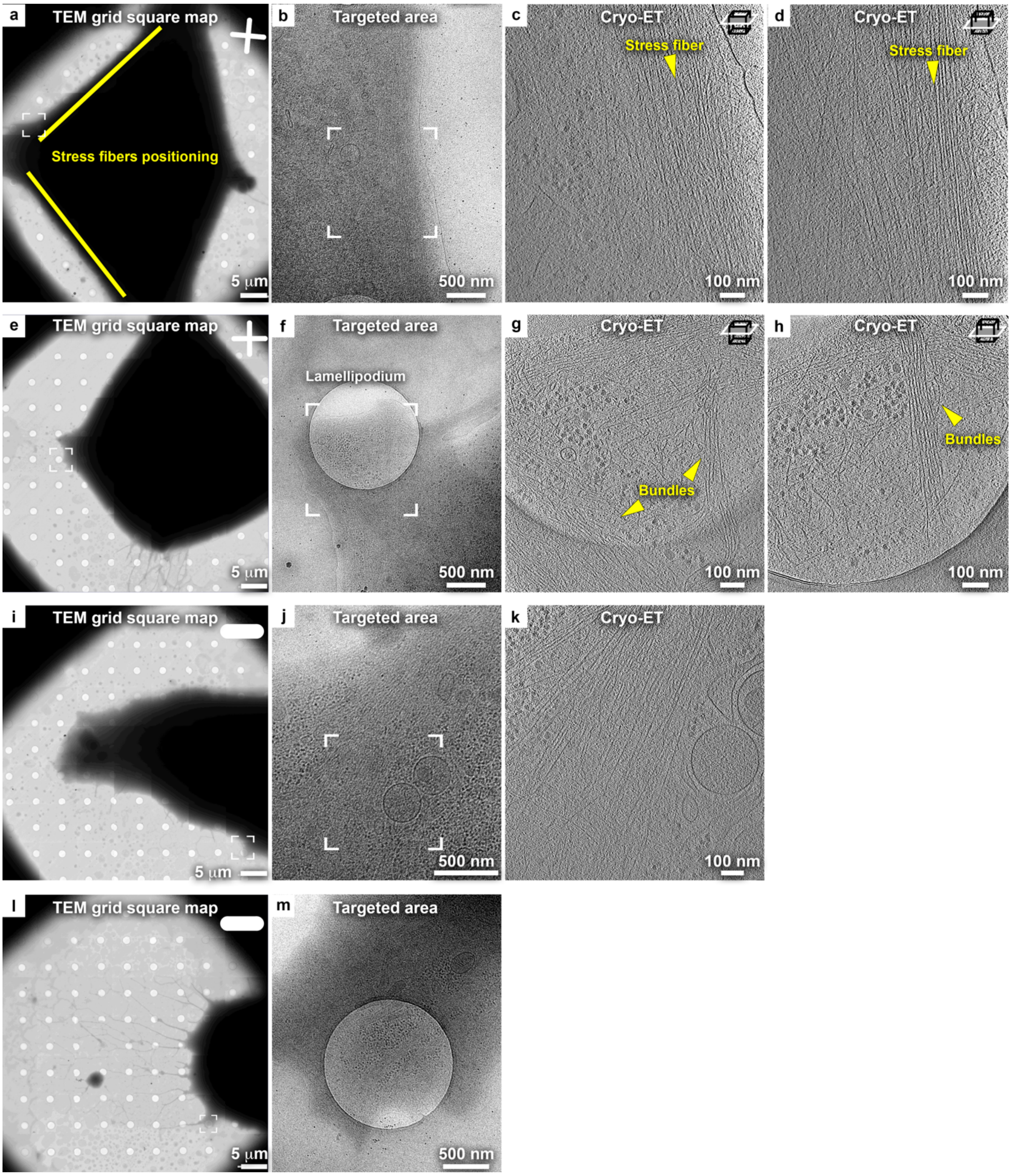


**Supplementary Figure 5. Direct cryo-ET assessment of actin networks in RPE1 cells grown on various micropattern shapes.**

**(a)** Cryo-TEM micrograph of the grid square from Fig. 2c (grown on a cross-shaped pattern). Yellow lines indicate the expected positioning of peripheral stress fibers as indicated from live cell imaging of the LifeAct fluorescence (Fig. 2a, top left). **(b)** Cryo-TEM micrograph of the framed area in (a) (rotated 90° clockwise), displaying a stress fiber targeted for tomography. **(c-d)** Tomographic slices at two different *z*-positions (6.8 nm thickness) through the 3D volume of the specified area in (b), showing the organization of actin filaments into a stress fiber. **(e)** Cryo-TEM micrograph of a different RPE1 cell grown on a cross-shaped pattern from the grid shown in Fig. 2b. **(f)** Cryo-TEM micrograph of the framed area in (e) (rotated 90° clockwise), displaying a lamellipodium with intricate actin architecture. **(g-h)** Tomographic slices at two different *z*-positions (6.8 nm thickness) through the 3D volume of the specified area in (f), showing the organization of actin filaments into multiple interrelated bundles. **(i)** Cryo-TEM micrograph of an RPE1 cell grown on an oval-shaped pattern from the grid shown in Fig. 2b. **(j)** Cryo-TEM micrograph of the framed area in (i) (rotated 90° clockwise), displaying the targeted area. **(k)** Tomographic slice (6.8 nm thickness) of the periphery of the cell depicting actin filaments bundling. **(l)** Cryo-TEM micrograph of RPE1 cell related to Fig. 2f, spreading on an oval-shaped pattern. **(m)** Cryo-TEM micrograph of the framed area in (l), displaying the targeted area represented in Fig. 2f.

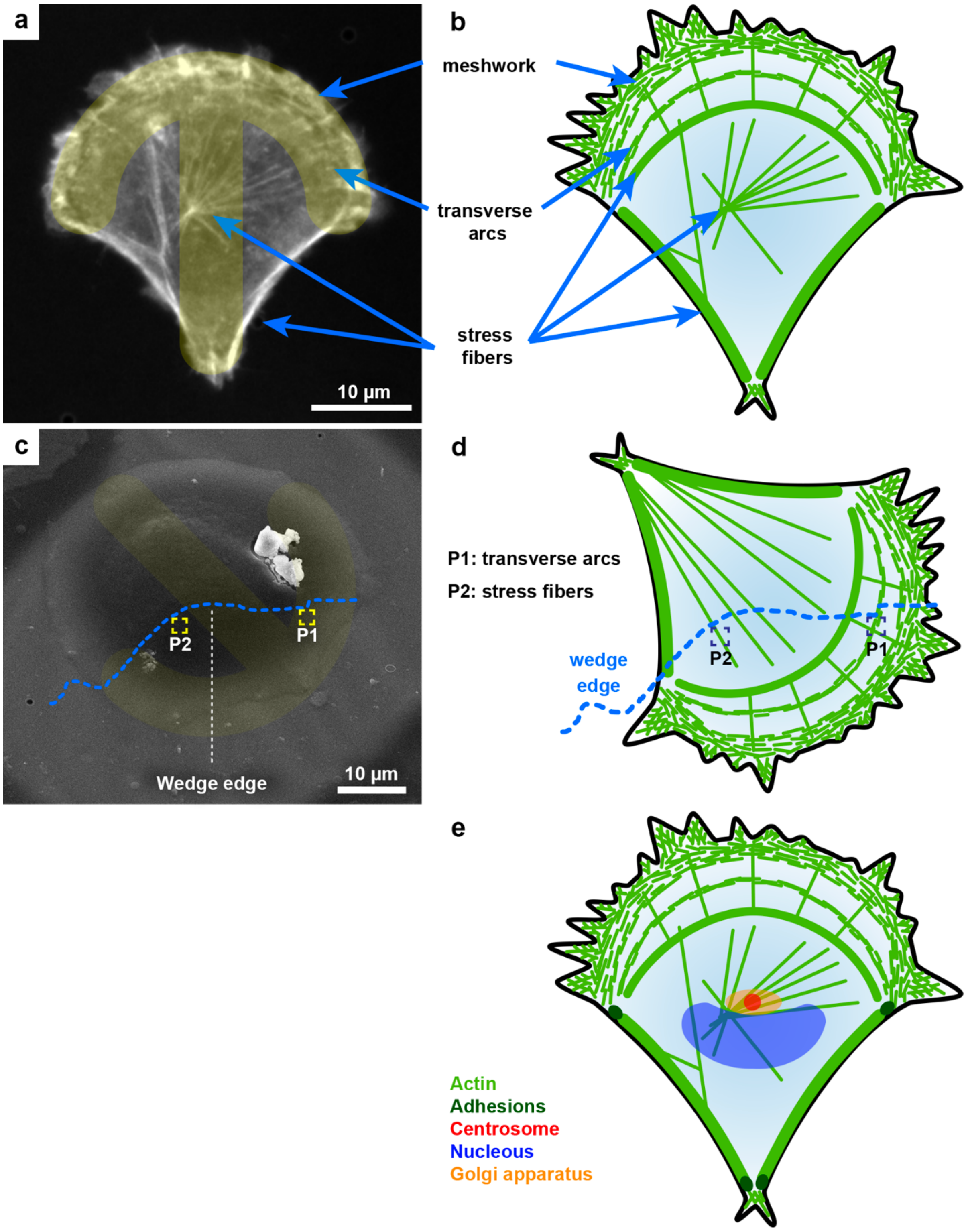

**Supplementary Figure 6. Actin networks architecture in RPE1 LifeAct-GFP cells grown on crossbow pattern.**

**(a)** On-grid live-cell Airyscan confocal slice at the basal region of an RPE1 cell (shown in Fig. 2a, top right) grown on a crossbow fibronectin-coated micropattern, and overlaid with the pattern design (yellow). **(b)** Actin network organization throughout the whole cell correlates with physical cues provided by cell-shape restriction of the micropattern shape (in agreement with[13]): (i) an extended actin meshwork in the cellular periphery (at the edge of the arc) forms a lamellipodium in adhesive regions of the pattern. This network extends towards the inner part of the cell with (ii) perpendicular fibers and parallel arcs of filament bundles. (iii) Stress fibers connect regions of the cell positioned on non-adhesive passivated areas of the support**. (c-d)** Correlation of actin network organization throughout the whole cell to the cryo-ET data acquired on a cryo-FIB generated wedge from Fig. 2h. **(c)** SEM top view image of an RPE1 cell grown on a crossbow fibronectin-coated micropattern. A wedge was produced to allow access to the inner basal part of the cell (dashed blue line corresponds to the edge of the wedge SEM image on the original micromachined cell). **(d)** Schematic representation of the expected actin map of an RPE1 cell on a crossbow micropattern overlapped with the location of the wedge edge and tomograms positioning. **(e)** Schematic representation of the actin architecture as well as organelles positioning of RPE1 cells spread on a crossbow-shaped micropattern in accordance with[13]. Models are not to scale.

### *Supplementary Video Legends*

**Supplementary Video 1.** Time-lapse, Live-cell imaging of HeLa cells expressing GFP-tagged β-tubulin (Cyan) and mCherry-tagged histone (H2B-mCherry: magenta) 8 h post-release from an S-phase cell-cycle block. Fields of view of a single grid square of a non-patterned (left) and micropatterned grid (right) are shown. Related to Fig. 1c-d. Gold-grids with a $SiO_2$ (R1/20) holey film. Scale: 20 µm.

**Supplementary Video 2.** Tomographic volume of the periphery of an RPE1 cell grown on a cross-shape pattern. Conventional defocus tomogram denoised with an anisotropic nonlinear diffusion algorithm. Video related to Fig. 2e. Each slice is 6.8 nm thick. Scale: 100 nm.

**Supplementary Video 3.** Tomographic volume of the periphery of an RPE1 cell grown on an oval-shape pattern. Conventional defocus tomogram. Video related to Fig. 2f. Each slice is 6.8 nm thick. Scale: 100 nm.

**Supplementary Video 4.** Tomographic volume of a cryo-FIB generated wedge of an RPE1 cell grown on a crossbow-shape pattern. VPP defocus tomogram Video related to Fig. 2j, wedge position 1. Each slice is 6.8 nm thick. Scale: 100 nm.
